# The Complete Chloroplast Genome of *Dendrobium nobile*, an endangered medicinal orchid from Northeast India and its comparison with related chloroplast genomes of *Dendrobium* species

**DOI:** 10.1101/316778

**Authors:** Ruchishree Konhar, Manish Debnath, Atanu Bhattacharjee, Durai Sundar, Pramod Tandon, Debasis Dash, Devendra Kumar Biswal

## Abstract

The medicinal orchid genus *Dendrobium* belonging to the Orchidaceae family is the largest genus comprising about 800-1500 species. To better illustrate the species status in the genus *Dendrobium*, a comparative analysis of 33 newly sequenced chloroplast genomes retrieved from NCBI Refseq database was compared with that of the first complete chloroplast genome of *D. nobile* from north-east India based on next-generation sequencing methods (Illumina HiSeq 2500-PE150). Our results provide comparative chloroplast genomic information for taxonomical identification, alignment-free phylogenomic inference and other statistical features of *Dendrobium* plastomes, which can also provide valuable information on their mutational events and sequence divergence.

## Introduction

*Dendrobium* is a huge genus of the tribe Dendrobieae (Orchidaceae: Epidendroideae) that was established by Olof Swartz in 1799. The present-day *Dendrobium* is the largest genus with approximately 800-1500 species and occurs in diverse habitats throughout much of Southeast Asia, including China, Japan, India, and the Philippines, Indonesia, New Guinea, Vietnam, Australia and many of the islands in the Pacific. [1].

Many species and cultivars of this genus are well-known floral motifs and have featured in artwork. *Dendrobium* orchids are popular not only for their visual appeal in cut flower market, but also for their herbal medicinal history of about 2000 years in east and south Asian countries [2]. Due to their diverse medicinal values namely for treating kidney and lung ailments, as a potent tonic for treating gastrointestinal problems and strengthening body’s immunity, improving sexual potency, anti-cancerous properties, treatment for lumbago and arthralgia etc., many species in this genus have been extensively used as herbal medicines for several hundreds of years. However, many *Dendrobium* species in the wild face an extreme threat of extinction due to their low germination and slow growth rate, habitat decline and over exploitation arising out of anthropogenic activities [3].

*Dendrobium* orchids have overwhelmed researchers because of their high economic importance in global horticultural trade and in Asian traditional medicine leading to extensive plant systemic studies particularly in species identification, novel marker development, breeding and conservation. In the past two decades, promising advances have been made in areas of molecular taxonomy and systematics and selective breeding of *Dendrobium* species by intensive use of molecular markers. Recently, a variety of molecular markers like microsatellite (SSR), Random amplified polymorphic DNA (RAPD) and amplified fragment length polymorphism (AFLP) markers including several other DNA barcode markers from different loci of nuclear and chloroplast (cp) regions have been developed to study *Dendrobium* diversity. However, these species are notoriously difficult to identify [4].

The complete cp genome usually contains a uniparentally inherited DNA, a feature which makes it an obvious choice for plant taxonomical analyses, phylogenomics and phylogeographic inferences at different taxonomic levels. Studies pertaining to plastome genome sequences are useful in investigating the maternal inheritance in plants, especially those with polyploid species, owing to their high gene content and conserved genome structure [5,6 & 7]. The advent of High-throughput sequencing technologies has enabled a rapid increase in the rate of completion of cp genomes with faster and cheaper methods to sequence organellar genomes [8, 9]. At the time of writing this manuscript, chloroplast genomes from 33 *Dendrobium* species have been reported as per NCBI Organellar genome records (https://www.ncbi.nlm.nih.gov/genome/browse#!/organelles/dendrobium).

Of the many highly prized medicinal plants in the genus *Dendrobium*, *D. nobile* Lindl. is one such endangered medicinal orchid listed in the Convention on International Trade in Endangered Species of Wild Fauna and Flora (CITES) Appendix II that demands immediate attention for its protection and propagation. Here, we report the first complete chloroplast genome of *D. nobile* from north-east India based on next-generation sequencing methods (Illumina HiSeq 2500-PE150) and further compare its structure, gene arrangement and microsatellite repeats with related species of 33 other newly sequenced *Dendrobium* chloroplast genomes. Our results provide comparative chloroplast genomic information for taxonomical identification, phylogenomic inference and other statistical features of *Dendrobium* plastomes, which can also provide valuable information on their mutational events and sequence divergence. The availability of complete cp genome sequences of these species in the genus *Dendrobium* will benefit future phylogenetic analyses and aid in germplasm utilization of these plants.

## Materials and Methods

### Sample collection, DNA extraction and sequencing

Fresh leaves of *D. nobile* were collected from plants growing in greenhouse of National Research Centre for Orchids, Sikkim, India and voucher specimen was deposited with Botanical Survey of India as well as with the Department of Botany, North Eastern Hill University, Shillong. The high molecular weight DNA was extracted using a modified CTAB buffer, and treated according to a standard procedure for next generation sequencing on Illumina HiSeq 2500-PE150. The quality and quantity of the genomic DNA was assessed through agarose gel electrophoresis, Nanodrop and Qubit detection method. The experiments included both paired-end and mate-pair libraries. Tagmentation was carried out with ~4μg of Qubit quantified DNA and the tagmented sample was washed using AMPURE XP beads (Beckman Coulter #A63881) and further exposed to strand displacement. The strand-displaced sample of 2-5kb and 8-13kb gel was size selected and taken for overnight circularization. The linear DNA was digested using DNA Exonuclease. Subsequently the circularized DNA molecules were sheared using Covaris microTUBE, S220 system (Covaris, Inc., Woburn, MA, USA) for obtaining fragments in the range 300 to 1000bp. M280 Streptavidin beads (ThermoFisher Scientific, Waltham, MA) was used to cleanse the sheared DNA fragments with biotinylated junction adapters. Further the bead-DNA complex was subjected to End repair, A-Tailing and Adapter ligations.

### Data processing

The data quality assessment for Illumina WGS raw reads was carried out using FastQC tool. Perl scripts were written for adapter clipping and low quality filtering. Cp genomes of *D. officinale*, *D. huoshanense* and *D. strongylanthum* retrieved from NCBI-Refseq database was used as reference for the assembly. BWA-MEM algorithm with default parameter settings was used for aligning the adapter clipped and low quality trimmed processed reads with the *Dendrobium* cp genomes [10]. SPAdes-3.6.0 program was used for k-mer based (k-mer used 21, 33, 55 and 77) de-novo assembly with the aligned reads and the quality of the assembled genome was gauged using Samtools and Bcftools (read alignment and genome coverage calculation) [11]. (https://samtools.github.io/bcftools/bcftools.html).

### Genome annotation and codon usage

Basic Local Alignment Search Tool (BLAST; BLASTN, PHI-BLAST and BLASTX) [12], chloroplast genome analysis platform (CGAP) [13] and Dual Organellar GenoMe Annotator (DOGMA) [14] was used to annotate protein-coding and ribosomal RNA genes. The boundaries of each annotated gene with putative start, stop, and intron positions were manually determined by comparison with homologous genes from other orchid chloroplast genomes. Further tRNA genes were predicted using tRNAscan-SE [15] and ARAGORN [16]. RNA editing sites in the protein-coding genes of *D. nobile* were predicted using Plant RNA Editing Prediction & Analysis Computer Tool (PREPACT) (http://www.prepact.de). For this analysis, *D. nobile* cp genome was BLAST aligned against *Nicotiana tabacum*, *Oryza sativa*, *Japonica* Group, *Phalaenopsis aphrodite* subsp. *Formosana*, *Physcomitrella patens* subsp. Patens and *Zea mays* with a cutoff E-value set to 0.08. The circular genome maps were drawn in CLC Genomics Workbench followed by manual modification. The sequencing data and gene annotation were submitted to GenBank with accession number KX377961. MEGA 7 was used to analyze and calculate GC content, codon usage, nucleotide sequence statistics and relative synonymous codon usage (RSCU) [17].

### Simple sequence repeats analysis

MISA (http://pgrc.ipk-gatersleben.de/misa/misa.html), a tool for the identification and location of perfect microsatellites and compound microsatellites was used to search for potential simple sequence repeats (SSRs) loci in the cp genomes of all the *Dendrobium* species. The threshold point for SSRs identification was set to 10, 5, 4, 3, and 3 for mono-, di-, tri-, tetra-, and penta-nucleotides SSRs, respectively. All the repeats found were manually curated and the redundant ones were removed.

### Phylogenetic reconstruction with whole genome alignment and rearrangement analysis

For phylogenetic reconstruction we included *D. nobile* cp genomes from India and China along with 32 other *Dendrobium* cp genomes retrieved from GenBank. Four *Goodyera* species were taken as outgroup. The cp genome sequences were aligned with MAFFT v7.0.0 [18] and manually curated by visual inspection. The complete cp genome sequences and protein coding genes (PCGs) were used for the Bayesian phylogenetic reconstruction using MRBAYES 3.2.6 [19]. To further validate our results we employed “K-mer Based Tree Construction" in CLC Genomics Workbench that uses single sequences or sequence lists as input and creates a distance-based phylogenetic tree. For visualization and testing the presence of genome rearrangement and inversions, gene synteny was performed using MAUVE as implemented in DNASTAR 12.3 with default settings. Comparative analysis of nucleotide diversity (*Pi*) among the complete cp genomes of *Dendrobium* was performed using MEGA 7.

### Single Nucleotide Polymorphism identification and phylogenetic analysis without genome alignment

Phylogenetic tree was constructed based on the Single Nucleotide Polymorphisms (SNPs) identified in the whole chloroplast genomes using kSNP3.0 with default settings except for k-mer size [20]. SNPs were identified with k-mer size set to 23 based on which approximately 79% of the k-mers generated from median-length genome were unique.

## Results and Discussion

### Genome organization and features

The complete cp genome of *D. nobile* was determined from the data generated out of a whole genome project initiative of the same species by Paired-end and Mate pair data from Illumina HighSeq with 150*2 and Illumina NextSeq500 with 75*2 respectively. Further the aligned Illumina reads were separated and assembled using CLC Main Workbench Version 7.7.1 into the single longest scaffold. The *D. nobile* cp genome is a typical circular double-stranded DNA with a quadripartite structure; it is 152,018 bp in size and consists of Large Single Copy (LSC) (1..84,944; 84,944 bp), Small Single Copy (SSC) (111,230..125,733; 14504 bp), and two Inverted Repeat (IR) regions of 26,285 bp: IRA (84,945..111,229) and IRB (125,734..152018). In total 134 unique genes (79 PCGs, 8 rRNA genes, 7 pseudogenes and 38 tRNA genes) were successfully annotated, of which 12 genes {rps16, atpF, rpoC1, ycf3, rps12 (2), clpP, petB, rpl2 (2), ndhB (2)} are reported with introns (Fig. 1). We could identify a total of 20, 81 and 11 genes duplicated in the IR, LSC and SSC regions respectively in the *D. nobile* cp genome.

**Figure 1.**
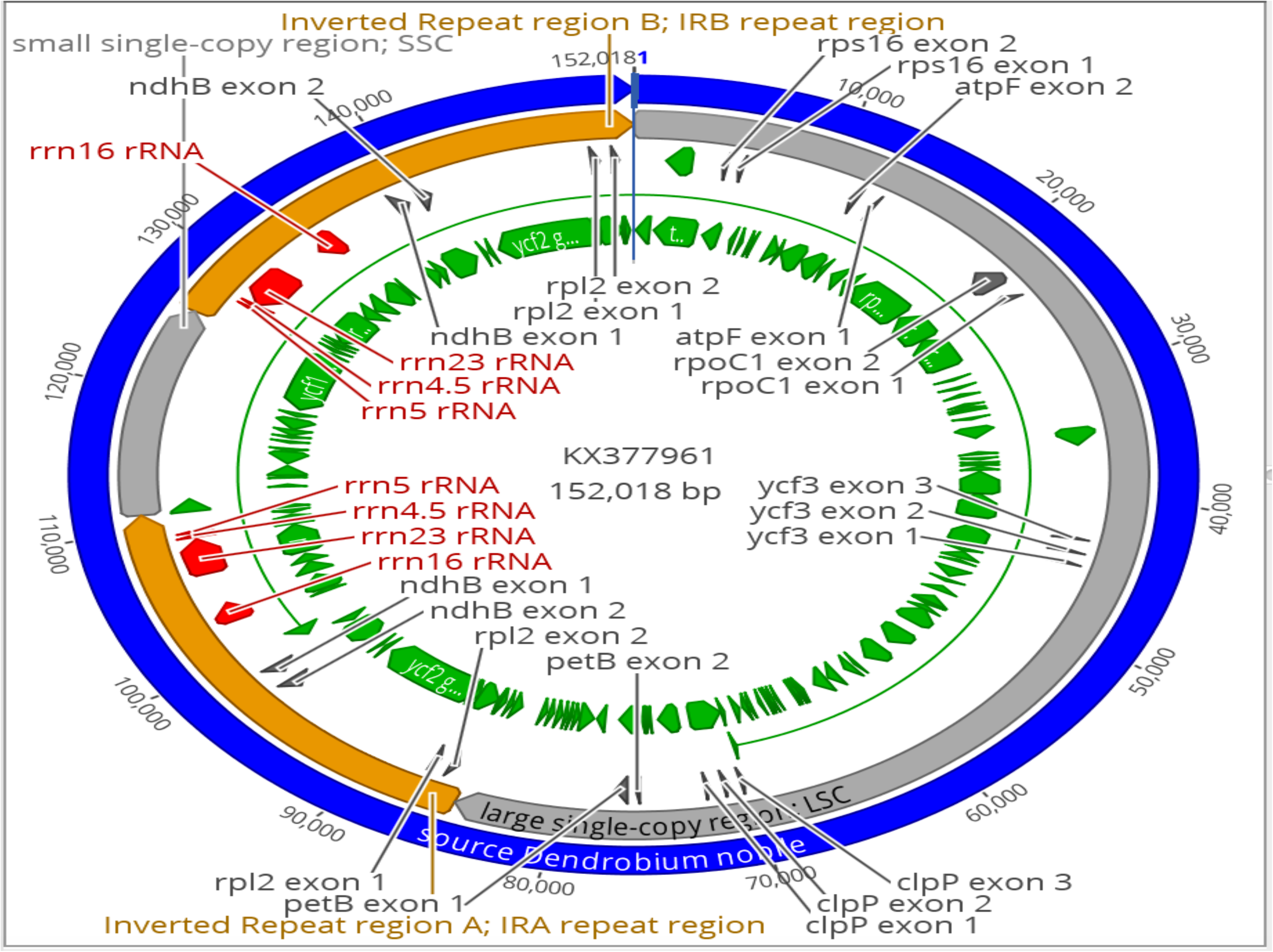
Gene map of *Dendrobium nobile* chloroplast genome from Northeast India. Genes shown inside the circle are transcribed clockwise, and those outside are transcribed anticlockwise. Color coding indicates genes of different functional groups. A pair of inverted repeats (IRA and IRB) separate the genome into LSC and SSC regions.

### Potential RNA editing sites

All 49 RNA editing sites (Table 1) were congruently predicted in 23 genes of *D. nobile* from at least 75% of the reference organisms (*Nicotiana tabacum*, *Oryza sativa Japonica* Group, *Phalaenopsis aphrodite* subsp. *Formosana*, *Physcomitrella patens* subsp. Patens and *Zea mays*) and resulted in amino acid substitutions.. All the RNA-editing sites were non-silent and edited C to U. Of the 49 RNA editing sites 89.8% (44) editing sites appeared in the second position of triplet codon, 10.2% (5) editing sites appeared in the first position of triplet codon whereas no editing sites appeared in the third base of triplet codon. The genes ndhD, rpoB, rpoC1 had 8, 6 and 4 RNA editing sites respectively. All the 49 RNA editing sites led to changes in the amino acid. The most frequent amino acid conversion was hydrophilic to hydrophobic (S to L, 22 occurrences and S to F, 8 occurrences), followed by hydrophobic to hydrophobic conversions (P to L, 12 occurrences). Seven conversions were found to be hydrophilic to hydrophilic (H to Y, 5 occurrences and T to M, 2 occurrences).

**Table 1.** RNA editing sites predicted in *Dendrobium nobile* chloroplast genome

| Gene | Nucleotide Position | Amino Acid Position | Triplet position within codon | Bases | Codon change | Amino Acid Conversion | Count | Percentage |
| --- | --- | --- | --- | --- | --- | --- | --- | --- |
| matK | 1258 | 420 | 1 | C→U | CAC→UAC | H→Y | 4/5 | 80% |
|  | 913 | 305 | 1 | C→U | CAU→UAU | H→Y | 4/6 | 80% |
| rps16 | 143 | 48 | 2 | C→U | UCA→UUA | S→L | 4/7 | 100% |
| atpA | 773 | 258 | 2 | C→U | UCA→UUA | S→L | 4/8 | 100% |
| atpF | 92 | 31 | 2 | C→U | CCA→CUA | P→L | 4/9 | 100% |
| atpI | 629 | 210 | 2 | C→U | UCA→UUA | S→L | 4/10 | 100% |
|  | 428 | 143 | 2 | C→U | CCU→CUU | P→L | 4/11 | 100% |
| rpoC1 | 617 | 206 | 2 | C→U | UCG→UUG | S→L | 4/12 | 100% |
|  | 488 | 163 | 2 | C→U | UCA→UUA | S→L | 4/13 | 100% |
|  | 182 | 61 | 2 | C→U | UCU→UUU | S→F | 4/14 | 100% |
|  | 41 | 14 | 2 | C→U | CCA→CUA | P→L | 4/15 | 100% |
| rpoB | 2426 | 809 | 2 | C→U | UCA→UUA | S→L | 4/16 | 80% |
|  | 623 | 208 | 2 | C→U | CCG→CUG | P→L | 4/17 | 80% |
|  | 566 | 189 | 2 | C→U | UCG→UUG | S→L | 4/18 | 100% |
|  | 551 | 184 | 2 | C→U | UCA→UUA | S→L | 4/19 | 100% |
|  | 473 | 158 | 2 | C→U | UCG→UUG | S→L | 4/20 | 100% |
|  | 338 | 113 | 2 | C→U | UCU→UUU | S→F | 4/21 | 100% |
| rps14 | 149 | 50 | 2 | C→U | CCA→CUA | P→L | 4/22 | 100% |
| ycf3 | 191 | 64 | 2 | C→U | CCA→CUA | P→L | 4/23 | 100% |
|  | 185 | 62 | 2 | C→U | ACG→AUG | T→M | 4/24 | 100% |
|  | 44 | 15 | 2 | C→U | UCU→UUU | S→F | 4/25 | 100% |
| atpB | 1184 | 395 | 2 | C→U | UCA→UUA | S→L | 4/26 | 100% |
| accD | 1184 | 395 | 2 | C→U | UCA→UUA | S→L | 4/27 | 100% |
|  | 1412 | 471 | 2 | C→U | CCA→CUA | P→L | 4/28 | 100% |
|  | 1430 | 477 | 2 | C→U | CCU→CUU | P→L | 4/29 | 100% |
| psaI | 80 | 27 | 2 | C→U | UCU→UUU | S→F | 4/30 | 100% |
| psbF | 77 | 26 | 2 | C→U | UCU→UUU | S→F | 4/31 | 100% |
| petL | 5 | 2 | 2 | C→U | CCU→CUU | P→L | 4/32 | 100% |
| rpl20 | 308 | 103 | 2 | C→U | UCA→UUA | S→L | 4/33 | 80% |
| clpP | 559 | 187 | 1 | C→U | CAU→UAU | H→Y | 4/34 | 100% |
|  | 82 | 28 | 1 | C→U | CAU→UAU | H→Y | 4/35 | 100% |
| petB | 611 | 204 | 2 | C→U | UCA→UUA | S→L | 4/36 | 100% |
| rpoA | 830 | 277 | 2 | C→U | UCA→UUA | S→L | 4/37 | 100% |
|  | 368 | 123 | 2 | C→U | UCA→UUA | S→L | 4/38 | 100% |
|  | 200 | 67 | 2 | C→U | UCU→UUU | S→F | 4/39 | 75% |
| rpl2 | 2 | 1 | 2 | C→U | ACG→AUG | T→M | 4/40 | 100% |
| ndhD | 878 | 293 | 2 | C→U | UCA→UUA | S→L | 4/41 | 100% |
|  | 674 | 225 | 2 | C→U | UCG→UUG | S→L | 4/42 | 100% |
|  | 383 | 128 | 2 | C→U | UCA→UUA | S→L | 4/43 | 100% |
| ndhA | 473 | 158 | 2 | C→U | UCA→UUA | S→L | 4/44 | 100% |
| ndhB | 149 | 50 | 2 | C→U | UCA→UUA | S→L | 4/45 | 100% |
|  | 467 | 156 | 2 | C→U | CCA→CUA | P→L | 4/46 | 100% |
|  | 586 | 196 | 1 | C→U | CAU→UAU | H→Y | 4/47 | 100% |
|  | 704 | 235 | 2 | C→U | UCC→UUC | S→F | 4/48 | 100% |
|  | 737 | 246 | 2 | C→U | CCA→CUA | P→L | 4/49 | 100% |
|  | 830 | 277 | 2 | C→U | UCA→UUA | S→L | 4/50 | 80% |
|  | 836 | 279 | 2 | C→U | UCA→UUA | S→L | 4/51 | 80% |
|  | 1481 | 494 | 2 | C→U | CCA→CUA | P→L | 4/52 | 100% |
| rpl23 | 71 | 24 | 2 | C→U | UCU→UUU | S→F | 4/53 | 80% |

### Comparison with other chloroplast genomes within the genus *Dendrobium*

We compared thirty-four chloroplast genomes representing different species within the genus *Dendrobium* (Table 2). The length of the *Dendrobium* species cp genomes ranged from 148,778 to 153,953 bp, with *D. chrysotoxum* being the largest cp genome and *D. moniliforme* the smallest. The cp genomes have acquired the familial angiosperm plastome organization comprising of a LSC, an SSC and a pair of IR regions each. *Dendrobium* cp genomes are also AT-rich (62.26–62.39%) quite similar to other orchid cp genomes. Differences in the cp genome size of these species are primarily due to the variations in the length of LSC, SSC and IR regions. Synteny comparison revealed a lack of genome rearrangement and inversions, thereby, substantiating for the highly conserved nature in the genomic structure, including gene number and gene order in these cp genomes. However, structural variation was predominant in the LSC/IR/SSC boundaries (Fig. 2), which could be harnessed for predicting potential biomarkers for species identification.

**Figure 2.**
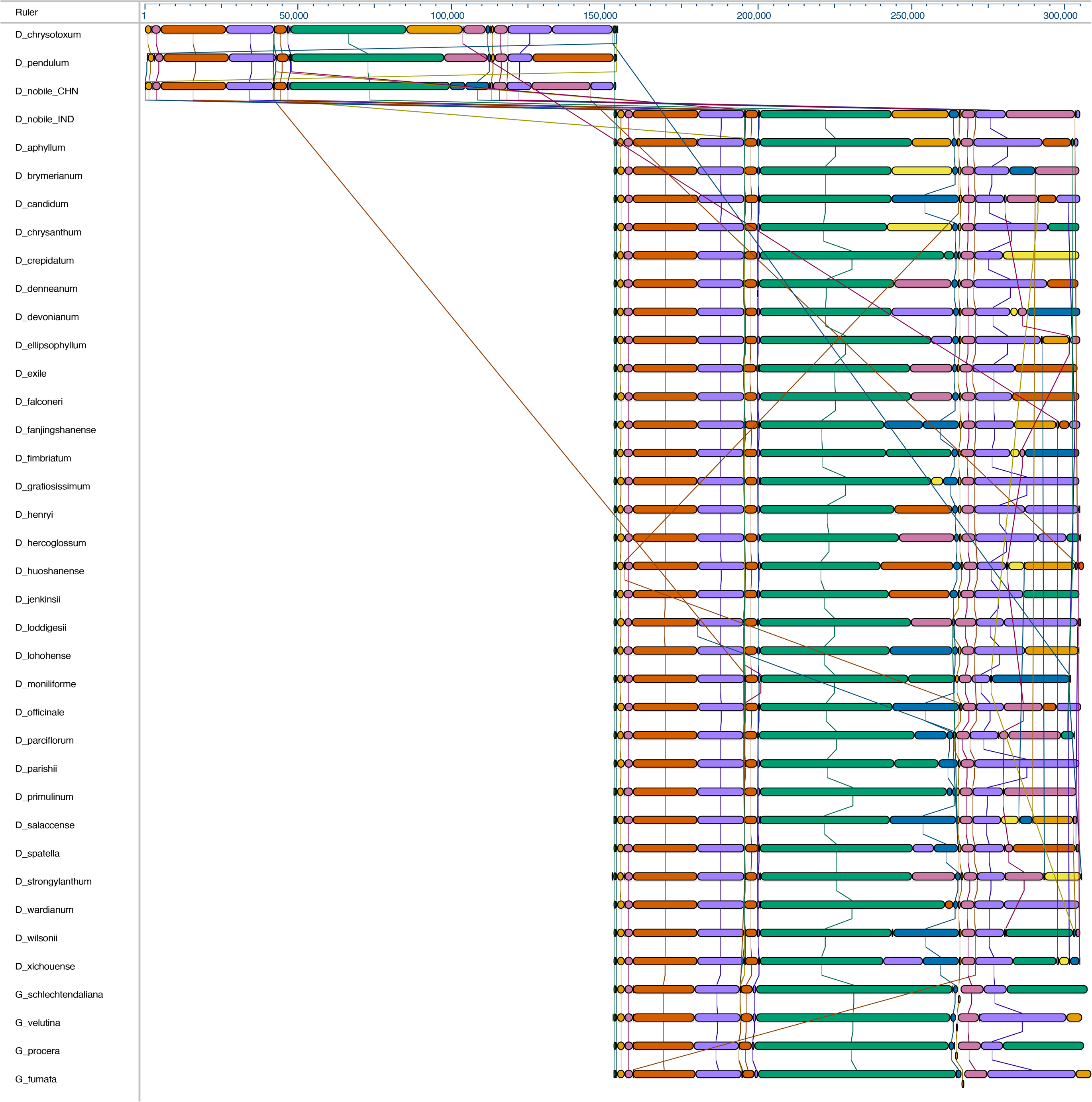
Whole chloroplast genome alignment of 38 orchid species. The whole chloroplast genome alignment includes 34 Dendrobium species and 4 species from the genus Goodyera as outgroup. Each genome’s panel contains its name, sequence coordinates and a black coloured horizontal centre line with coloured block outlines appearing above and below it. Each block represents homology with the genes, internally free from genomic rearrangement, connected by lines to similarly coloured blocks depicting comparative homology across genomes.

**Table 2.** Summary of characteristics in chloroplast genome sequences of thirty-four *Dendrobium* species and four *Goodyera* species (taken as outgroup).

| Organism | Accession Number | Length | Weight (single-stranded) Mda | Weight (double-stranded) Mda |
| --- | --- | --- | --- | --- |
| *Dendrobium nobile* | KX377961 | 152018 | 46.932 | 93.912 |
| *Dendrobium officinale* | NC_024019 | 152221 | 46.995 | 94.038 |
| *Dendrobium strongylanthum* | NC_027691 | 153059 | 47.256 | 94.556 |
| *Dendrobium huoshanense* | NC_028430 | 153188 | 47.294 | 94.635 |
| *Dendrobium chrysotoxum* | NC_028549 | 153953 | 47.528 | 95.108 |
| *Dendrobium nobile (China)* | NC_029456 | 153660 | 47.453 | 94.927 |
| *Dendrobium pendulum* | NC_029705 | 153038 | 47.246 | 94.542 |
| *Dendrobium moniliforme* | NC_035154 | 148778 | 45.931 | 91.911 |
| *Dendrobium primulinum* | NC_035321 | 150767 | 46.545 | 93.14 |
| *Dendrobium aphyllum* | NC_035322 | 151524 | 46.779 | 93.607 |
| *Dendrobium brymerianum* | NC_035323 | 151830 | 46.873 | 93.796 |
| *Dendrobium denneanum* | NC_035324 | 151565 | 46.793 | 93.633 |
| *Dendrobium devonianum* | NC_035325 | 151945 | 46.909 | 93.867 |
| *Dendrobium falconeri* | NC_035326 | 151890 | 46.891 | 93.833 |
| *Dendrobium gratiosissimum* | NC_035327 | 151829 | 46.873 | 93.796 |
| *Dendrobium hercoglossum* | NC_035328 | 151939 | 46.908 | 93.864 |
| *Dendrobium wardianum* | NC_035329 | 151788 | 46.861 | 93.77 |
| *Dendrobium wilsonii* | NC_035330 | 152080 | 46.951 | 93.951 |
| *Dendrobium crepidatum* | NC_035331 | 151717 | 46.837 | 93.726 |
| *Dendrobium salaccense* | NC_035332 | 151104 | 46.648 | 93.347 |
| *Dendrobium spatella* | NC_035333 | 151829 | 46.872 | 93.796 |
| *Dendrobium parviflorum* | NC_035334 | 150073 | 46.331 | 92.711 |
| *Dendrobium henryi* | NC_035335 | 151850 | 46.88 | 93.809 |
| *Dendrobium chrysanthum* | NC_035336 | 151790 | 46.861 | 93.772 |
| *Dendrobium jenkinsii* | NC_035337 | 151717 | 46.839 | 93.726 |
| *Dendrobium lohohense* | NC_035338 | 151812 | 46.868 | 93.785 |
| *Dendrobium parishii* | NC_035339 | 151689 | 46.83 | 93.709 |
| *Dendrobium ellipsophyllum* | NC_035340 | 152026 | 46.935 | 93.917 |
| *Dendrobium xichouense* | NC_035341 | 152052 | 46.942 | 93.933 |
| *Dendrobium fimbriatum* | NC_035342 | 151673 | 46.825 | 93.699 |
| *Dendrobium exile* | NC_035343 | 151294 | 46.707 | 93.465 |
| *Dendrobium fanjingshanense* | NC_035344 | 152108 | 46.96 | 93.968 |
| *Dendrobium candidum* | NC_035745 | 152094 | 46.955 | 93.959 |
| *Dendrobium loddigesii* | NC_036355 | 152493 | 47.077 | 94.205 |
| *Goodyera fumata* | NC_026773 | 155643 | 48.048 | 96.151 |
| *Goodyera procera* | NC_029363 | 153240 | 47.306 | 94.667 |
| *Goodyera schlechtendaliana* | NC_029364 | 154348 | 47.648 | 95.351 |
| *Goodyera velutina* | NC_029365 | 152692 | 47.138 | 94.328 |

IR regions are generally considered to be highly conserved regions in the chloroplast genome. In the evolutionary ladder, SSC and IR border regions experience expansion and contraction that overall contribute to the variation in chloroplast genome length [21, 22]. Thus, the positions of LSC/IRA/SSC/IRB borders were examined in the overall alignment of *Dendrobium* whole cp genomes and all of them were found to have similar structures at the IR/LSC junction (Fig. 3).

**Figure 3.**
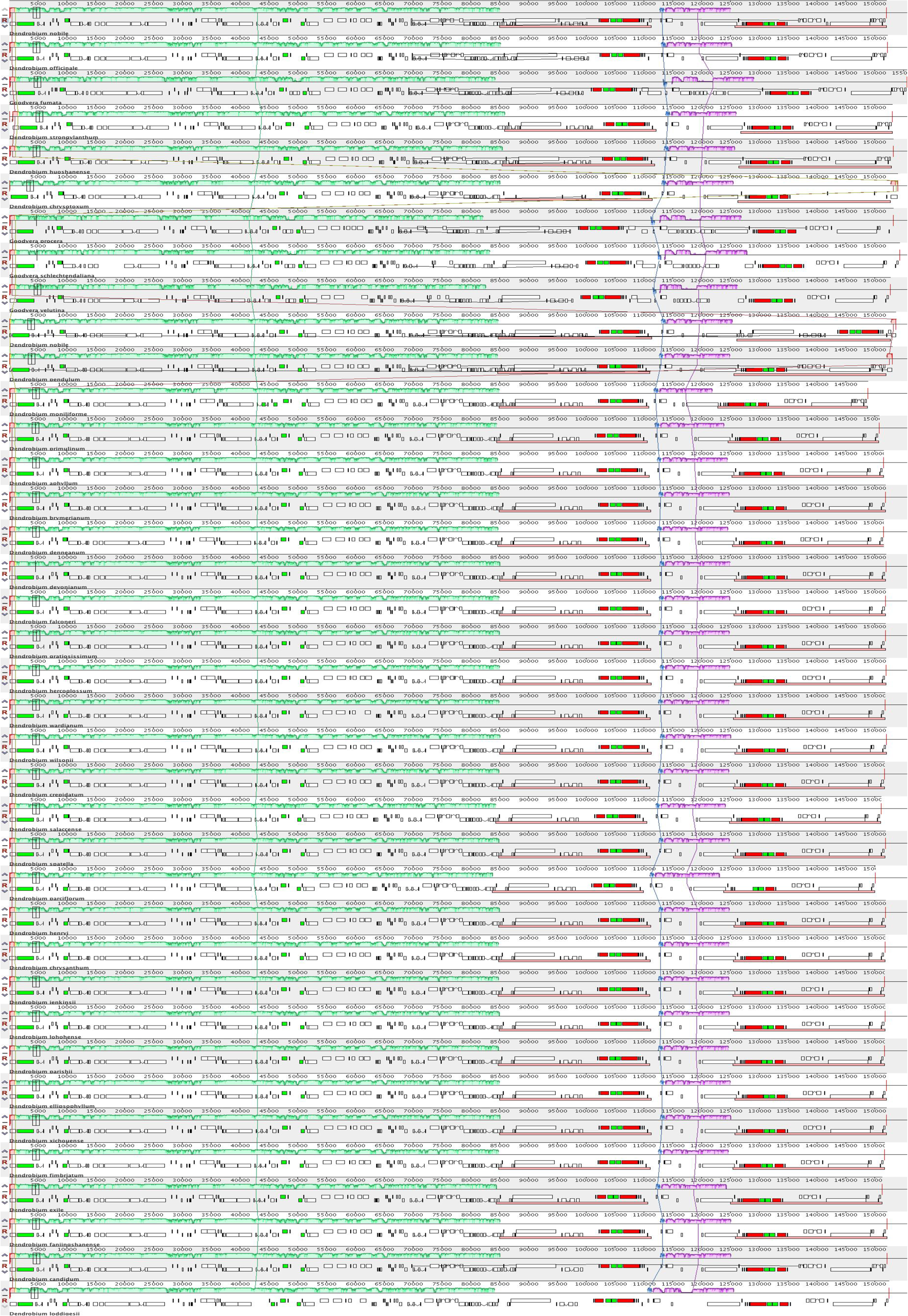
Comparison of the borders of LSC, SSC and IR regions across *Dendrobium*chloroplast genomes.

A comparative nucleotide sequence statistics (counts of annotations, AT/GC counts, nucleotide frequency in codon positions etc.) for all the *Dendrobium* species including representatives from outgroup are outlined in Tables 3, 4 and 5. The relative synonymous codon usage is given in parentheses following the codon frequency (averages over all taxa) involved (Table 6). Maximum Likelihood analysis of natural selection codon-by-codon was carried out. For each codon, estimates of the numbers of inferred synonymous (s) and nonsynonymous (n) substitutions are presented along with the number of sites that are estimated to be synonymous (S) and nonsynonymous (N) (Table S1). These estimates were calculated using the joint Maximum Likelihood reconstructions of ancestral states under a Muse-Gaut model [23] of codon substitution and Felsenstein 1981 model [24] of nucleotide substitution. For estimating ML values, a tree topology was automatically computed. The test statistic dN - dS was used for detecting codons that have undergone positive selection, where dS is the number of synonymous substitutions per site (s/S) and dN is the number of nonsynonymous substitutions per site (n/N). A positive value for the test statistic indicates an overabundance of nonsynonymous substitutions. In this case, the probability of rejecting the null hypothesis of neutral evolution (P-value) was calculated [25, 26]. Values of P less than are considered significant at a 5% level and are highlighted [Table S2]. Normalized dN - dS for the test statistic is obtained using the total number of substitutions in the tree (measured in expected substitutions per site). The analysis involved 38 nucleotide sequences. Codon positions included were 1st+2nd+3rd+Noncoding and all positions containing gaps and missing data were eliminated. There were a total of 108,594 positions in the final dataset.

**Table 3.**
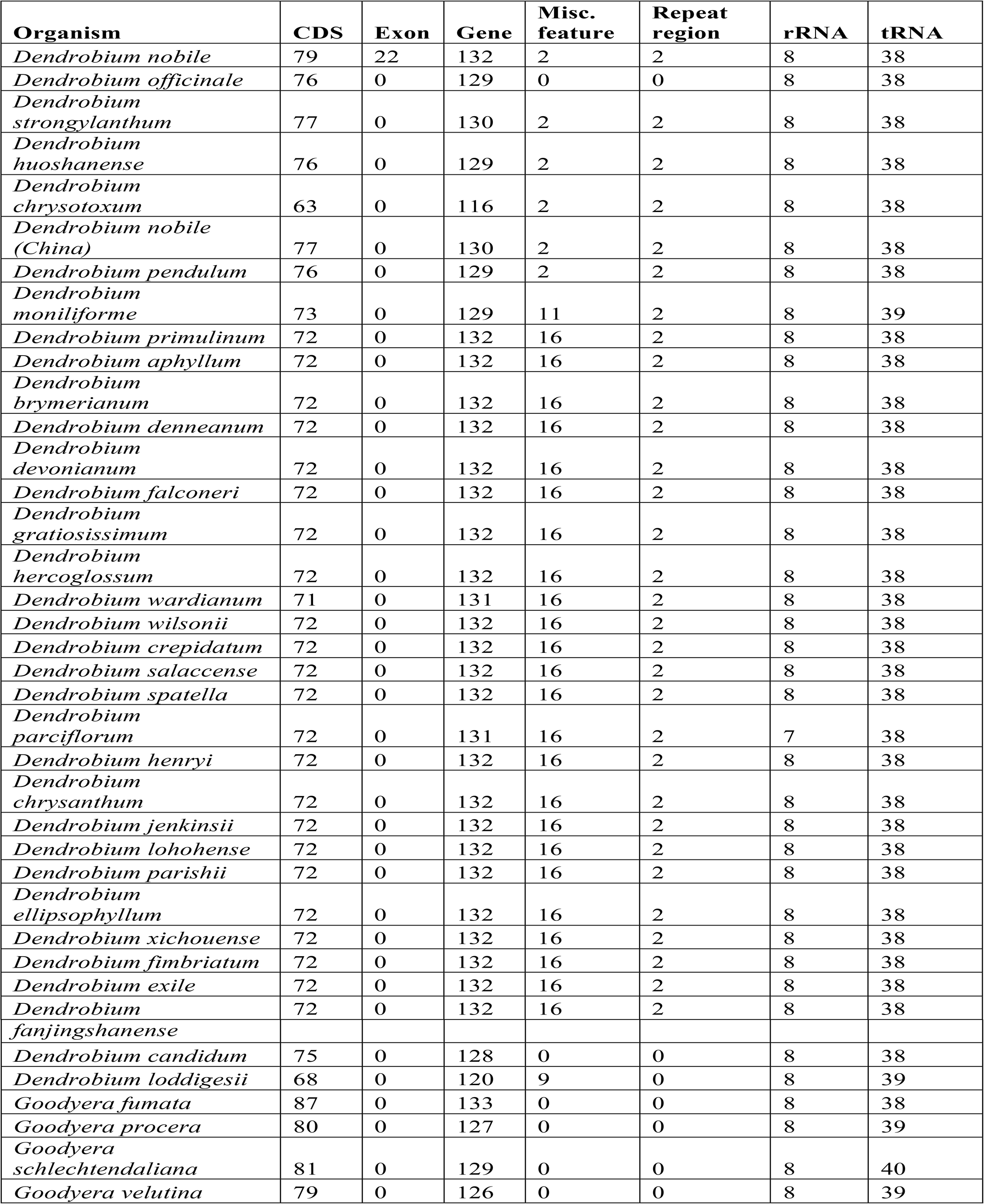
Summary features of chloroplast genome sequences of thirty-four *Dendrobium* species and four *Goodyera* species

**Table 4.** Counts of nucleotides in the chloroplast genomes.

| **Nucleotide** | **Adenine**<br>**(A)** | **Cytosine**<br>**(C)** | **Guanine**<br>**(G)** | **Thymine**<br>**(T)** | **C + G** | **A + T** |
| --- | --- | --- | --- | --- | --- | --- |
| *Dendrobium nobile* | 46576 | 28853 | 28039 | 48381 | 56892 | 94957 |
| *Dendrobium officinale* | 46743 | 28924 | 28107 | 48447 | 57031 | 95190 |
| *Dendrobium strongylanthum* | 46940 | 29147 | 28431 | 48541 | 57578 | 95481 |
| *Dendrobium huoshanense* | 47032 | 29111 | 28316 | 48729 | 57427 | 95761 |
| *Dendrobium chrysotoxum* | 47180 | 29400 | 28492 | 48881 | 57892 | 96061 |
| *Dendrobium nobile (China)* | 47118 | 28871 | 28748 | 48923 | 57619 | 96041 |
| *Dendrobium pendulum* | 46997 | 29122 | 28242 | 48677 | 57364 | 95674 |
| *Dendrobium moniliforme* | 45551 | 28339 | 27520 | 47368 | 55859 | 92919 |
| *Dendrobium primulinum* | 46191 | 28750 | 27909 | 47917 | 56659 | 94108 |
| *Dendrobium aphyllum* | 46417 | 28917 | 28057 | 48133 | 56974 | 94550 |
| *Dendrobium brymerianum* | 46509 | 28968 | 28123 | 48230 | 57091 | 94739 |
| *Dendrobium denneanum* | 46440 | 28913 | 28115 | 48097 | 57028 | 94537 |
| *Dendrobium devonianum* | 46615 | 28943 | 28108 | 48279 | 57051 | 94894 |
| *Dendrobium falconeri* | 46591 | 28911 | 28040 | 48348 | 56951 | 94939 |
| *Dendrobium gratiosissimum* | 46521 | 28954 | 28095 | 48259 | 57049 | 94780 |
| *Dendrobium hercoglossum* | 46592 | 28941 | 28131 | 48275 | 57072 | 94867 |
| *Dendrobium wardianum* | 46479 | 28955 | 28118 | 48236 | 57073 | 94715 |
| *Dendrobium wilsonii* | 46668 | 28948 | 28101 | 48363 | 57049 | 95031 |
| *Dendrobium crepidatum* | 46482 | 28951 | 28056 | 48228 | 57007 | 94710 |
| *Dendrobium salaccense* | 46493 | 28635 | 27735 | 48241 | 56370 | 94734 |
| *Dendrobium spatella* | 46524 | 28969 | 28091 | 48245 | 57060 | 94769 |
| *Dendrobium parciflorum* | 45941 | 28699 | 27829 | 47604 | 56528 | 93545 |
| *Dendrobium henryi* | 46550 | 28936 | 28093 | 48271 | 57029 | 94821 |
| *Dendrobium chrysanthum* | 46519 | 28939 | 28078 | 48254 | 57017 | 94773 |
| *Dendrobium jenkinsii* | 46497 | 28942 | 28105 | 48173 | 57047 | 94670 |
| *Dendrobium lohohense* | 46558 | 28928 | 28098 | 48228 | 57026 | 94786 |
| *Dendrobium parishii* | 46487 | 28924 | 28079 | 48199 | 57003 | 94686 |
| *Dendrobium ellipsophyllum* | 46690 | 28922 | 28091 | 48323 | 57013 | 95013 |
| *Dendrobium xichouense* | 46672 | 28937 | 28098 | 48345 | 57035 | 95017 |
| *Dendrobium fimbriatum* | 46483 | 28932 | 28094 | 48164 | 57026 | 94647 |
| *Dendrobium exile* | 46251 | 28937 | 28065 | 48041 | 57002 | 94292 |
| *Dendrobium fanjingshanense* | 46694 | 28947 | 28115 | 48352 | 57062 | 95046 |
| *Dendrobium candidum* | 46695 | 28914 | 28091 | 48394 | 57005 | 95089 |
| *Dendrobium loddigesii* | 46868 | 28934 | 28064 | 48627 | 56998 | 95495 |
| *Goodyera fumata* | 48186 | 29569 | 28447 | 49441 | 58016 | 97627 |
| *Goodyera procera* | 47095 | 29370 | 28303 | 48472 | 57673 | 95567 |
| *Goodyera schlechtendaliana* | 47822 | 29206 | 28146 | 49174 | 57352 | 96996 |
| *Goodyera velutina* | 47554 | 28694 | 27658 | 48786 | 56352 | 96340 |

**Table 5.**
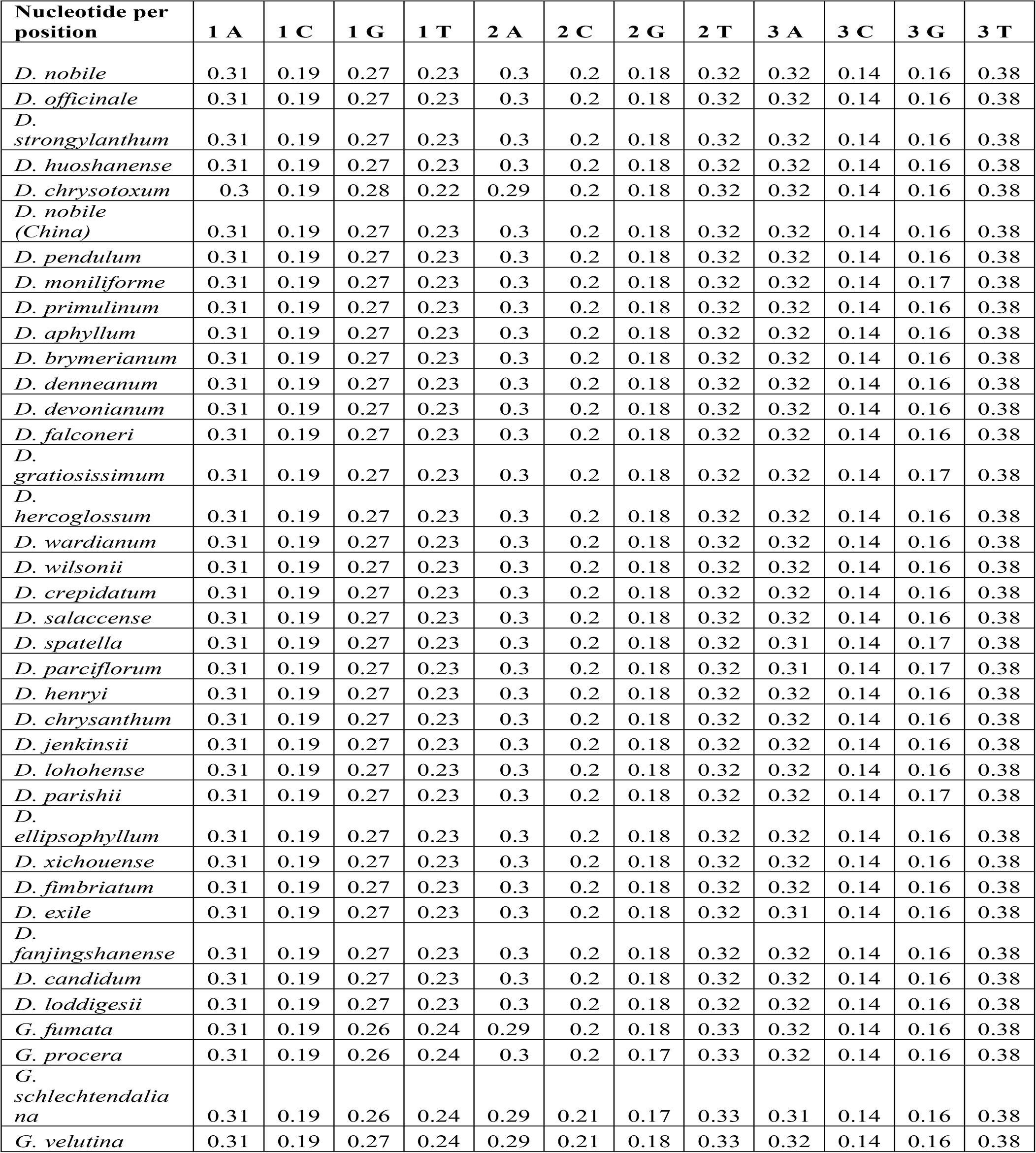
Counts of nucleotide frequency in codon positions across the chloroplast genomes.

**Table 6.** Relative synonymous codon usage (in parentheses) following the codon frequency across the chloroplast genomes in the genus *Dendrobium.*

| Codon | Count | RSCU | Codon | Count | RSCU | Codon | Count | RSCU | Codon | Count | RSCU |
| --- | --- | --- | --- | --- | --- | --- | --- | --- | --- | --- | --- |
| UUU(F) | 2018.1 | 1.16 | UCU(S) | 1330 | 1.63 | UAU(Y) | 1371 | 1.38 | UGU(C) | 706.9 | 1.24 |
| UUC(F) | 1459.2 | 0.84 | UCC(S) | 882.8 | 1.08 | UAC(Y) | 621.4 | 0.62 | UGC(C) | 437 | 0.76 |
| UUA(L) | 918.4 | 1.14 | UCA(S) | 999.4 | 1.23 | UAA(*) | 970.5 | 1.05 | UGA(*) | 1065 | 1.15 |
| UUG(L) | 970.9 | 1.21 | UCG(S) | 576.9 | 0.71 | UAG(*) | 732.2 | 0.79 | UGG(W) | 691.4 | 1 |
| CUU(L) | 1068.9 | 1.33 | CCU(P) | 638 | 1.13 | CAU(H) | 919.7 | 1.43 | CGU(R) | 336.1 | 0.63 |
| CUC(L) | 629.2 | 0.78 | CCC(P) | 547.8 | 0.97 | CAC(H) | 369.3 | 0.57 | CGC(R) | 220.7 | 0.41 |
| CUA(L) | 762.8 | 0.95 | CCA(P) | 689.4 | 1.23 | CAA(Q) | 952.8 | 1.38 | CGA(R) | 545.2 | 1.02 |
| CUG(L) | 473.7 | 0.59 | CCG(P) | 375.4 | 0.67 | CAG(Q) | 423.2 | 0.62 | CGG(R) | 343 | 0.64 |
| AUU(I) | 1635.7 | 1.21 | ACU(T) | 646 | 1.21 | AAU(N) | 1580 | 1.39 | AGU(S) | 659.9 | 0.81 |
| AUC(I) | 1072.9 | 0.8 | ACC(T) | 530.8 | 1 | AAC(N) | 695 | 0.61 | AGC(S) | 435.8 | 0.54 |
| AUA(I) | 1337.4 | 0.99 | ACA(T) | 610.3 | 1.15 | AAA(K) | 1914 | 1.31 | AGA(R) | 1171 | 2.2 |
| AUG(M) | 891.4 | 1 | ACG(T) | 343.2 | 0.64 | AAG(K) | 1009 | 0.69 | AGG(R) | 576 | 1.08 |
| GUU(V) | 709.4 | 1.36 | GCU(A) | 467.5 | 1.29 | GAU(D) | 1038 | 1.43 | GGU(G) | 523.7 | 0.99 |
| GUC(V) | 366.7 | 0.7 | GCC(A) | 326.4 | 0.9 | GAC(D) | 413.9 | 0.57 | GGC(G) | 314.4 | 0.59 |
| GUA(V) | 647.8 | 1.24 | GCA(A) | 438.7 | 1.21 | GAA(E) | 1335 | 1.37 | GGA(G) | 754.1 | 1.43 |
| GUG(V) | 366.9 | 0.7 | GCG(A) | 221.5 | 0.61 | GAG(E) | 618.3 | 0.63 | GGG(G) | 521.8 | 0.99 |

### Characterization of simple sequence repeats

SSRs were identified in MISA perl scripts with a minimum of 10 bp repeats among all the *Dendrobium* species. Of all the SSRs, the mononucleotide A/T repeat units occupied the highest proportion. A higher proportion of di-, tri-repeats are reported rather than tetra- and penta-nucleotide repeats across *Dendrobium* cp genomes (Fig. 4). The SSRs could be further investigated for identifying potential markers that can aid in barcoding analysis.

**Figure 4.**
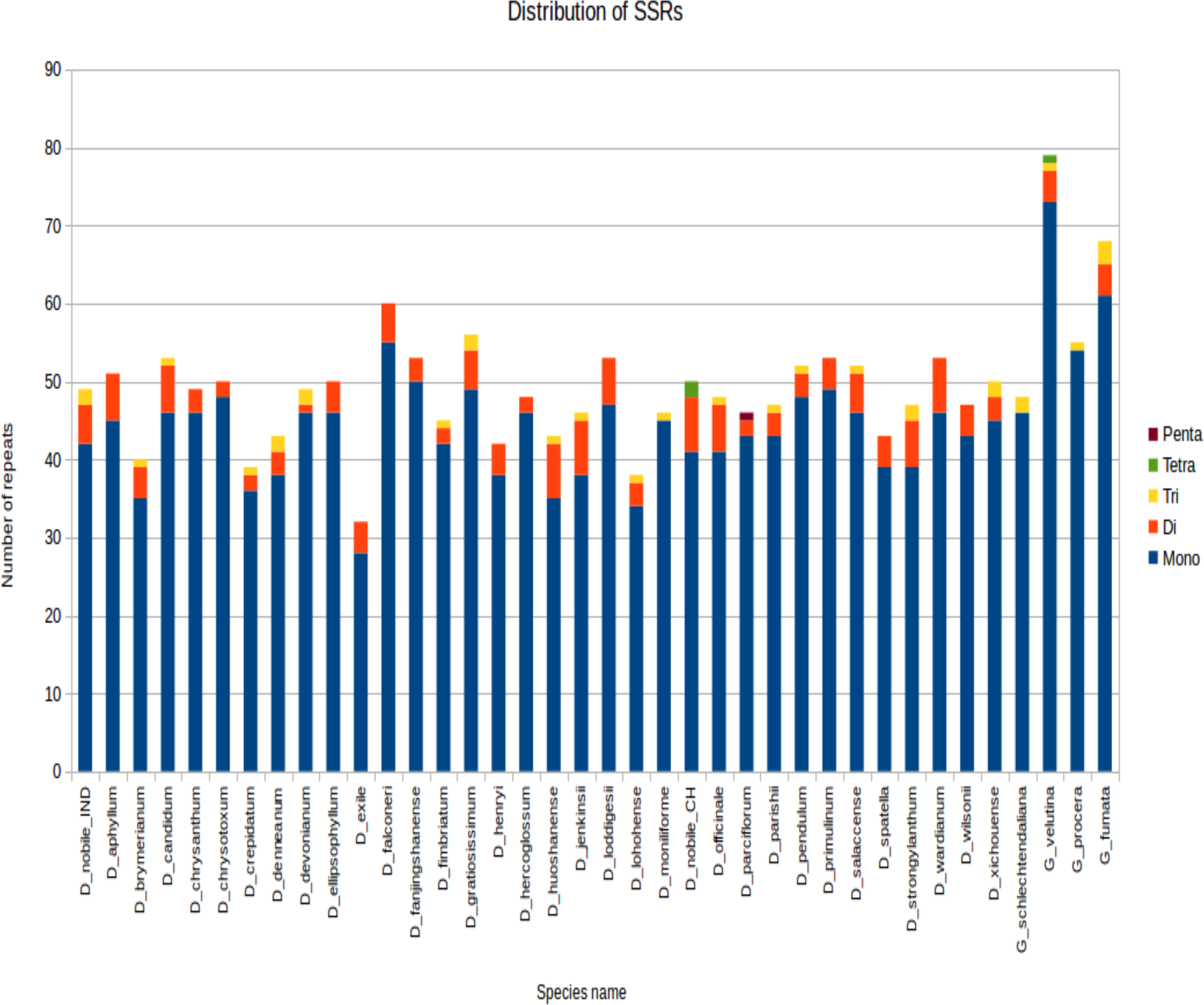
SSR distribution among different *Dendrobium* plastomes. The SSR were determined in MISA per scripts based on the comparison between plastomes of each tested *Dendrobium* species and *D*. *nobile*. Histograms with different color codes indicate the numbers of SSRs. The minimum number (thresholds) of SSRs was set as 10, 5, 4, 3, and 3 for mono-, di-, tri-, tetra-, and penta-nucleotides SSRs, respectively.

### Phylogenetic analyses

In the present study, we employed two different approaches for phylogeny reconstruction. First we aligned the whole cp genomes and exported the alignment matrices for creating a Bayesian tree (Fig. 5). Two independent MCMC chains were run with first 25% of the cycles removed as burn-in, coalescence of substitution rate and rate model parameters were also examined and average standard deviation of split frequencies was carried out and generations added until the standard deviation value was lowered to 0.01. Secondly we performed a phylogenetic tree construction using an alignment free approach. In this case we identified the SNPs from the cp genomes and utilised them in constructing the phylogenetic tree (Fig. 6). A total of 13,839 SNPs were identified in the 38 genomes analyzed, of which 2,203 were homoplastic SNPs i.e. SNPs that do no correspond to any node in the parsimony tree. The fraction of k-mers present in all genomes is 0.482. The numbers at the nodes in the phylogenetic tree indicate the number of SNPs that are present in all of the descendants of that node and absent in others (Fig. 6). The numbers at the tips indicate the number of SNPs unique to each particular species.

**Figure 5.**
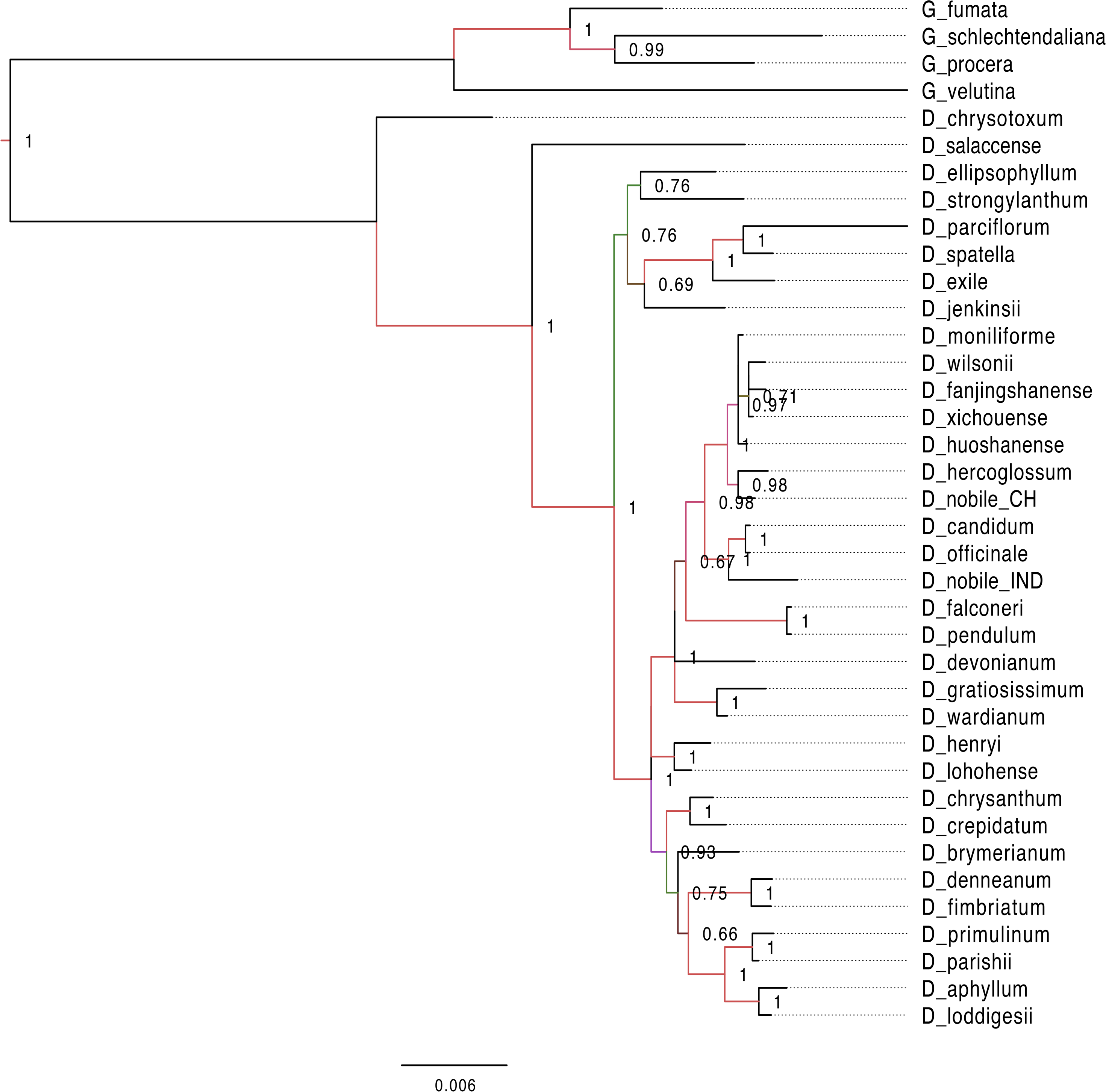
Phylogenetic tree based on bayesian inference from the whole genome alignment matrix of *Dendrobium* chloroplast genomes. The tree yielded monophyletic groupings of the genus *Dendrobium* and Goodyera species emerged as outgroup with a separate clade. Posterior probability/bootstrap values are indicated on the internal nodes, which are highly supportive of the overall tree topology.

**Figure 6.**
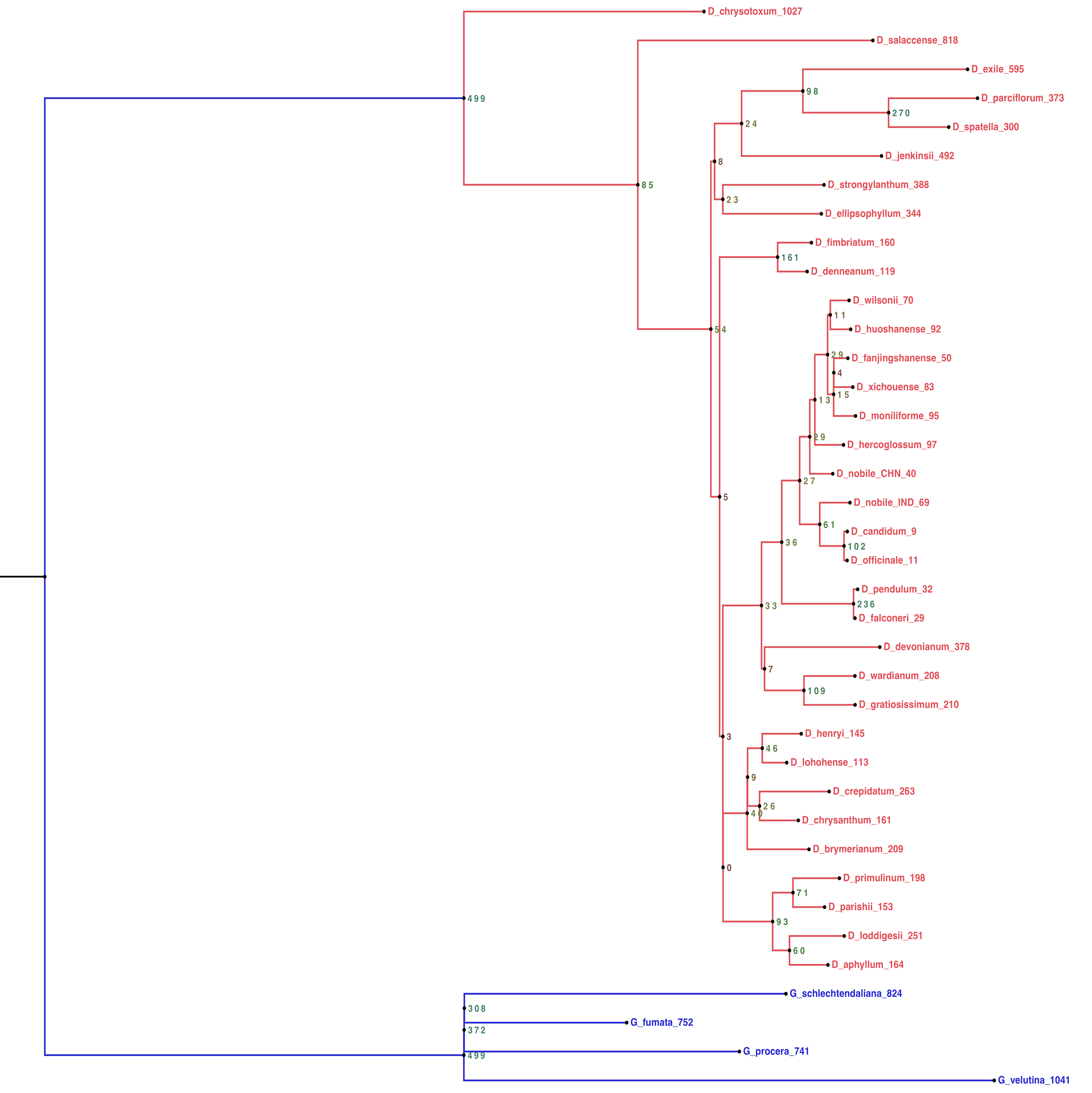
Alignment free phylogenetic tree reconstruction based on SNP identification. The optimum kmer size for the dataset is determined that calculates FCK, a measure of diversity of sequences in the dataset (Kchooser) and a consensus of the equally most parsimonious trees are reported. The numbers at the nodes indicate the number of SNPs that are present in all of the descendants of that node and absent in others. The numbers within parentheses at the tips indicate the number of SNPs unique to each particular species.

The two different methods that employed both alignment and alignment-free approach resulted in highly reliable identical phylogenetic trees within each data set. Different analyses based on the two datasets generated largely congruent topologies (Figs. 5 and 6) with *Dendrobium* species forming one clade and *Goodyera* species forming another clade as an outgroup.

## Conclusions

This study provides the first comparative account on the complete chloroplast genome of *D. nobile* from north-east India with 33 other species from the genus *Dendrobium* that revealed higher sequence variation in SSC and LSC regions compared with IR regions in both coding and non-coding regions. The gene order, gene content and genomic structure were highly conserved with those of other sequenced *Dendrobium* species. However, IR contraction is observed within the genus and several SNPs identified from these cp genomes were quite instrumental in generating alignment-free robust phylogenetic trees that congrued with trees generated from aligned matrices of whole cp genomes. This gives an indication that the SNPs, insertions and deletions, LSC and SSC regions in the cp genomes of this medicinal orchid genus can be utilized for barcoding and biodiversity studies. Further this would augment more and more plastome sequencing of *Dendrobium* species that are not reported in this study.

## Data Availability

The following information was supplied regarding the GenBank accession, BioSample, SRA and BioProject pertaining to this study.

NCBI GenBank accession number: KX377961.

BioSample: SAMN05190527; Sample name: SO_5373; SRA: SRS1473719 BioProject Accession: PRJNA323854; ID: 323854

## Competing Interests

The authors have declared that no competing interests exist.

## Author Contributions

Ruchishree Konhar performed the experiments, analyzed the data, contributed analysis tools, prepared figures and/or tables, authored and reviewed drafts of the paper. Manish Debnath performed the in silico analysis and prepared tables and figures. Atanu Bhattacharjee and Durai Sundar analyzed the data, contributed analysis tools, authored or reviewed drafts of the paper, and approved the final draft. Debasis Dash provided physical resources, contributed analysis tools, authored or reviewed drafts of the paper, and approved the final draft. Devendra K Biswal and Pramod Tandon conceived and guided on the overall design of experiments, provided physical resources, contributed analysis tools, authored and reviewed drafts of the paper, approved the final draft.

## Acknowledgement

This work was funded under the DBT-NER Twinning program titled “Next Generation Sequencing (NGS)-based de novo assembly of expressed transcripts and genome information of Orchids in North-East India " (Grant ID BT/325/NE/TBP/2012 dated August 07, 2014) by the Department of Biotechnology (DBT), Government of India. RK is supported by University Grants Commission (UGC) Junior Research Fellowship. The funders had no role in study, design or preparation of the manuscript. We also thank Mr. Santosh Vishwakarma, Bioinformatics Centre for providing us with the list of SSRs from the *Dendrobium* species chloroplast genomes.

## Supplementary Files

**Table S1.** Maximum Likelihood analysis of natural selection codon-by-codon

**Table S2.** Results from the Fisher’s Exact Test of Neutrality Selection across the chloroplast genome sequences in the genus Dendrobium

